# Biosynthesis of mitis group streptococcal glycolipids and their roles in physiology and antibiotic susceptibility

**DOI:** 10.1101/2024.10.30.621112

**Authors:** Yahan Wei, Guan H. Chen, Muneer Yaqub, Elice Kim, Lily E Tillett, Luke R. Joyce, Nicholas Dillon, Kelli L. Palmer, Ziqiang Guan

**Affiliations:** School of Podiatric Medicine, The University of Texas Rio Grande Valley, Harlingen, Texas, USA; Department of Biological Sciences, The University of Texas at Dallas, Richardson, Texas, USA; Department of Immunology and Microbiology, University of Colorado Anschutz Medical Campus, Aurora, Colorado, USA; Department of Biochemistry, Duke University School of Medicine, Durham, North Carolina, USA

**Author notes:** **Corresponding authors (email)**: Yahan Wei, Ziqiang Guan.

## Abstract

Bacterial cell surface components such as lipoteichoic acids (LTAs) play critical roles in host-microbe interactions and alter host responses based on their chemical structures. Mitis group streptococci have commensal and pathogenic interactions with the human host and produce Type IV LTAs that are slightly different in chemical structures between species. To reveal the molecular bases for the intricate interactions between MGS and human hosts, a detailed understanding of the structure and biosynthetic process of MGS LTAs is needed. In this study, we used genomic and lipidomic techniques to elucidate the biosynthetic processes of Type IV LTA and its associated glycolipid anchors, monohexosyl-diacylglycerol and dihexosyl-diacyglycerol, in the infectious endocarditis isolate *Streptococcus* sp. strain 1643. Through establishing a murine sepsis model, we validated the essentiality of these glycolipids in the full virulence of *S. mitis*. Additionally, we found that these glycolipids play an important role in protecting the bacteria from antimicrobials. Overall, results obtained through this study both confirm and dispute aspects of the existing model of glycolipids biosynthesis, provide insights into the fundamental roles of bacterial glycolipids, as well as suggest the potential of targeting glycolipids for developing antimicrobial therapeutics.

**Significance/Importance:** Glycolipids and glycolipid-anchored LTAs are common and essential membrane components in Gram-positive bacteria. Yet, the biosynthesis and functions of LTAs have not been fully understood for many significant Gram-positive pathogens. Through genomic, lipidomic, and animal infection model analyses, as well as antimicrobial susceptibility assays, this study advances our understanding of Type IV LTA biosynthesis and the physiological roles of glycolipids in mitis group streptococci. Overall, our work establishes the essentiality of glycolipids in both bacterial virulence and defense against antimicrobials.

## Introduction

The Mitis group streptococci (MGS) encompass more than 20 *Streptococcus* species that are primarily isolated from the human oral cavity and upper respiratory tract (1). Major members of MGS include the significant human pathogen *Streptococcus pneumoniae* and pioneer oral colonizers *S. mitis* and *S. oralis*. *S. pneumoniae* causes an array of serious infections including pneumonia and meningitis (2), attributes to ∼ 1 million annual deaths of children < 5 years old (3), and is one of the leading causes of death associated with antimicrobial resistance worldwide (4). Conversely, while *S. mitis* and *S. oralis* are among the leading causes of life-threatening diseases including bacteremia and infective endocarditis (IE) (5), they have complex interactions with the human host, providing both beneficial and detrimental effects (6–9). Understanding the factors that contribute to the discrepancies in microbe-host interactions of different MGS species will not only reveal further details of their pathogenesis but also shed light on pathogen-specific methods of preventing opportunistic infections.

Bacterial cell surface factors directly interact with host cells and modulate immune responses. One of the main immune-stimulating cell-surface factors produced by Gram-positive bacteria is teichoic acid (TA), a polymer that can be either attached to a membrane lipid anchor (i.e. lipoteichoic acid, LTA) or peptidoglycan (i.e. wall teichoic acid, WTA) (10, 11). Based on the chemical structure of the polymer repeating unit, currently identified LTAs can be separated into six major types, Type I to VI, each of which elicits distinct immune reactions (11, 12). *S. pneumoniae*, *S. oralis*, and *S. mitis* all produce Type IV LTA, whose repeating unit consists of 2-acetamido-4-amino-2,4,6-trideoxy-D-galactose (AATGal), ribitolphosphate (RboP), N-acetyl-D-galactosamine (GalNAc), phosphocholine (ChoP), and hexoses (Fig. 1A), with slight differences in both the chemical structure modifications and residue compositions across different strains and species (13–17). These differences arise from variance in the presence/absence of biosynthetic genes (15, 18), despite the prediction of a conserved biosynthetic pathway from comparative genome analysis (Fig 1A) (13).

**Fig 1:**
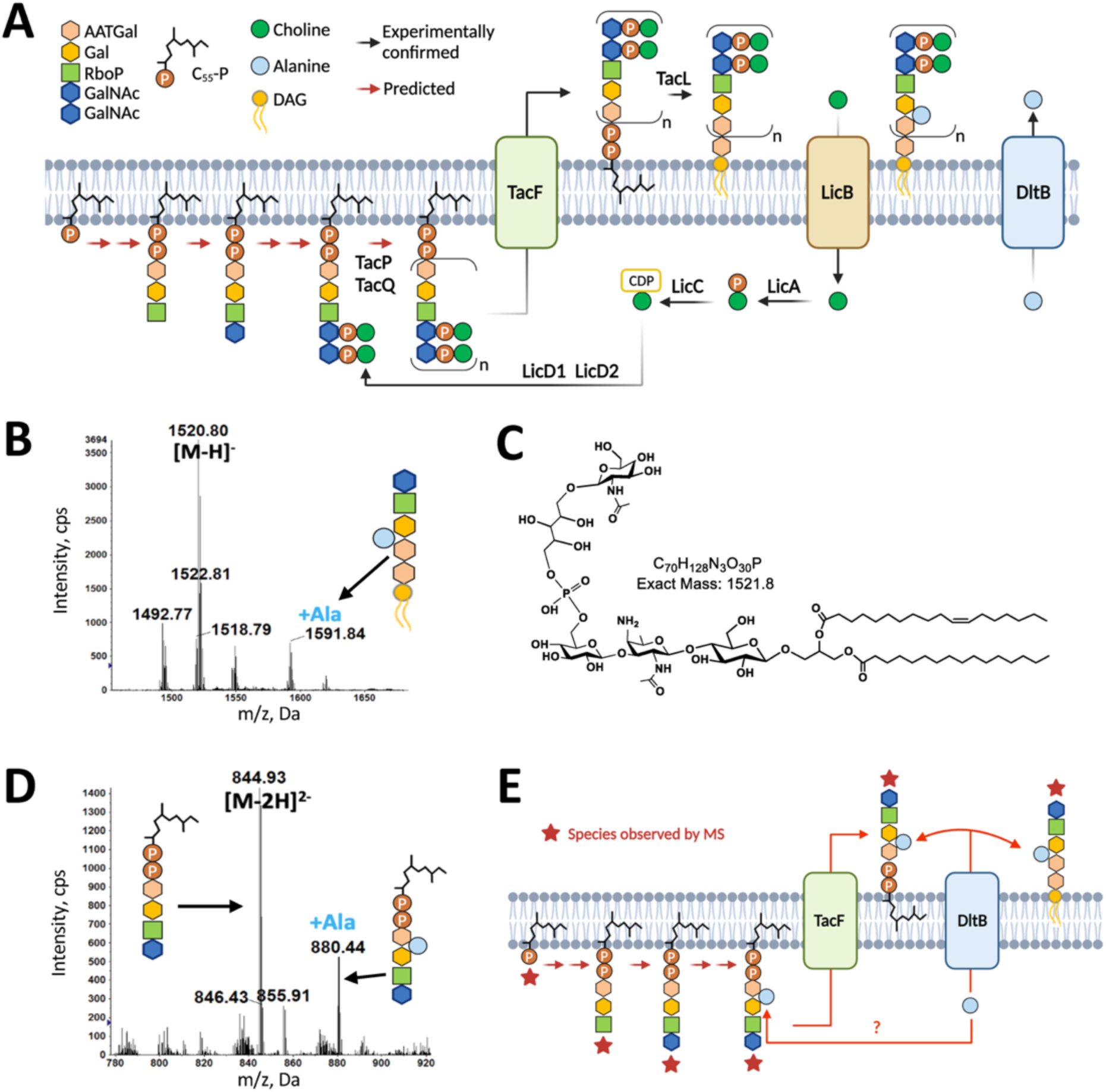
Detection of novel Type IV LTA biosynthetic intermediates. **A.** Diagram showing the proposed Type IV LTA biosynthetic process (13). Experimentally confirmed biosynthetic steps are indicated by black arrows (23, 51, 69, 70), while steps that have not been confirmed are indicated with red arrows. **B and D**. mass spectra (MS) of Type IV LTA intermediates detected in SM43 *ΔlicA* grown in TH broth and SM43 WT grown in CDM without choline: deprotonated [M-H]^-^ ions for GalNAc-RboP-Gal-AATGal-MHDAG (*m/z* 1520.8) and Ala-modified GalNAc-RboP-Gal-AATGal-MHDAG (*m/z* 1591.8) (B) and [M-2H]^2-^ ions for C_55_-PP-AATGal-Gal-RboP-GalNAc (*m/z* 844.9) and Ala-modified C_55_-PP-AATGal-Gal-RboP-GalNAc (*m/z* 880.4) (D). **C**. Chemical structure of GalNAc-RboP-Gal-AATGal-MHDAG. The depicted stereochemistry and linkage of hexose moieties are for illustration purposes and could not be determined by tandem MS. **E.** Alternative biosynthetic pathway of Type IV LTA in SM43 according to detected biosynthetic intermediates. Three biologically independent replicates were performed for each strain under the indicated conditions. The structural identification of C_55_-PP-linked pseudopentasaccharide and MHDAG-linked pseudopentasaccharide was further confirmed by MS/MS (Fig S2). **Abbreviations**: AATGal, 2-acetamido-4-amino-2,4,6-trideoxy-_D_-galactose; Gal, galactose; RboP, ribitol-phosphate; GalNAc, *N*-acetyl-_D_-galactosamine; C_55_-PP, undecaprenylpyrophosphate; MHDAG, monohexosyl-diacylglycerol.

Till now, LTA biosynthetic genes have only been validated in a few bacterial species that produce Type I LTA (19–21). The majority of the conserved genes of the Type IV LTA biosynthetic pathway have not been experimentally verified, mainly due to the essentiality of Type IV LTA, and specifically its ChoP residues, to *S. pneumoniae* growth (22) as well as the lack of sensitive analytical techniques required to monitor the LTA biosynthetic intermediates which are typically present at extremely low levels. LTAs serve as the docking sites for enzymes involved in cell replication (23, 24). In *S. pneumoniae*, LTA cannot be transported to the outer leaflet of the membrane without the proper incorporation of ChoP (25, 26); thus, lack of choline supplementation in growth media leads to the impairment of cell growth. Interestingly, *S. mitis* and *S. oralis* strains that do not require choline for proper growth have been identified (26–28), enabling experimental verification for the biosynthetic steps of Type IV in these strains. Additionally, it has been verified that *S. mitis* also produces Type I LTA (repeating unit is glycerophosphate (GroP)) (16), which potentially enables *S. mitis* to survive without producing Type IV LTA. In this study, we use normal-phase liquid chromatography (NPLC)-electrospray ionization/mass spectrometry (ESI/MS) to analyze the changes in membrane lipid compositions in the endocarditis isolate and MGS *Streptococcus sp.* strain 1643, referred to here as SM43 (29), grown with and without choline, providing experimental results that suggest an alternative biosynthetic trajectory for Type IV LTA than that proposed in prior literature (13, 15).

Moreover, we previously reported the detection of GroP-linked dihexosyl-diacylglycerol (DHDAG) in *S. oralis* and *S. pneumoniae*, strains that do not encode the Type I LTA synthase LtaS, and in *S. mitis* and *Staphylococcus aureus* that are deficient in producing LtaS (30). Additionally, structural analyses of glycolipids harvested from *S. mitis* identified two different monohexosyl-DAGs (MHDAGs), α-glucopyranosyl-(1,3)-DAG and β-galactofuranosyl-(1,3)-DAG), respectively serving as the lipid anchor of Type IV and Type I LTA (16). Results suggest the existence of novel GroP transferase(s), glycosyltransferase(s), and additional conserved functions of glycolipids in these bacteria. It is known that glycolipids participate in maintaining membrane curvature (31), serve as the lipid anchors for LTAs (32), and are involved in protection against cell surface targeting antimicrobials (33–35). In this study, we use genetically modified SM43 in conjunction with lipidomic analysis by LC/MS to confirm the functions of predicted glycosyltransferases in generating different species of glycolipids, i.e. DHDAG and MHDAG (these two glycolipids serve as the anchors for (Gro-P)-DHDAG and Type IV LTA respectively) and finally, the roles of these glycolipids in both bacterial physiology and virulence within an animal host.

## Results

### The endocarditis isolate SM43 can grow without choline, but becomes more susceptible to cell wall-targeting antibiotics

Based on the genetic compositions and amino acid similarities (Table S1), the Type IV LTA produced by SM43 has a higher similarity to that produced by *S. oralis* Uo5 rather than *S. pneumoniae*, which corresponds to the genomic analytical results indicating that SM43 is a strain closely related to *S. oralis* (36). Specifically, pneumococcal TA repeating unit polymerases TarP and TarQ are absent in SM43; and SM43 TacF shares higher amino acid similarity with *S. oralis* Uo5 TacF (>90%) rather than pneumococcal TacF (<50%). Considering that *S. oralis* has been reported as not requiring choline for optimum growth (37) and that pneumococcal choline dependency is determined by the sequence of TacF (25), the growth of SM43 may also be choline-independent. Indeed, SM43 can grow without the choline supplement (Fig 2A), though a significant growth defect compared to growth with choline is observed. As a comparison, *S. mitis* NCTC12261^T^ (referred to here as SM61), a strain with > 98% amino acid similarity in TacF with *S. pneumoniae*, barely grows without choline (Fig 2A). Additionally, SM43 grown without choline has increased susceptibility towards the cell wall-targeting antibiotics ampicillin and vancomycin (Fig 3A & B) compared to SM43 grown with the presence of choline, but no obvious change in susceptibility towards the protein synthesis inhibitor gentamicin (Fig 3C).

**Fig 2:**
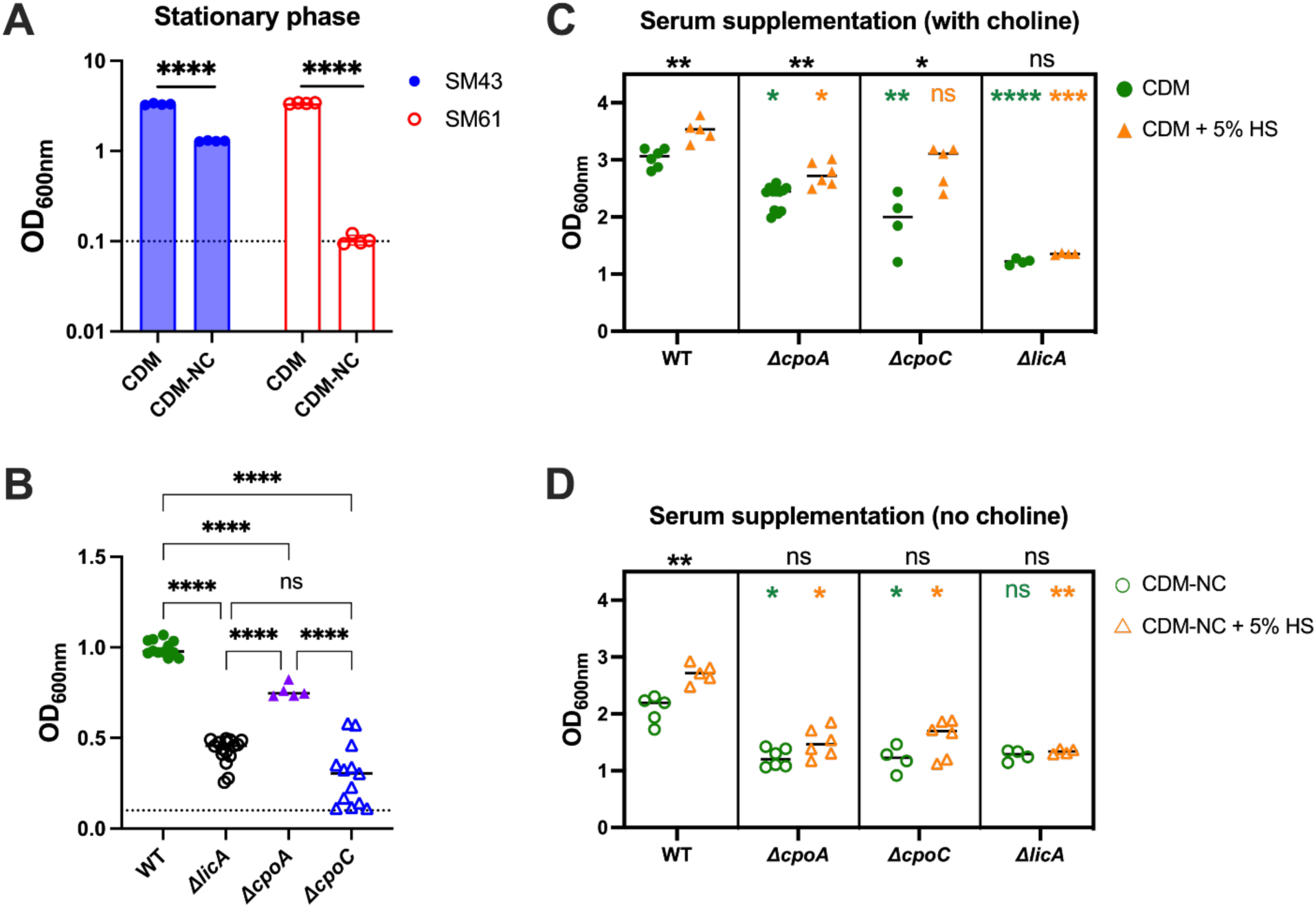
Stationary phase OD_600nm_ values of *S. mitis* NCTC12261^T^ (SM61), *Streptococcus* sp. 1643 (SM43) wildtype (WT), *ΔlicA*, *ΔcpoA*, and *ΔcpoC* strains. Bacterial single colonies were grown overnight in either choline-supplemented chemically defined medium (CDM) (A, C, & D) or Todd-Hewitt broth (THB) (B). For A, single-colony cultures were diluted to 0.1 of OD_600nm_ with either CDM or chemically defined medium without choline (CDM-NC) and grown overnight followed by another dilution to 0.1 of OD_600nm_ with the same conditions. OD_600nm_ values of the cultures were measured and plotted after overnight incubation. For B, C, & D, single-colony cultures were diluted to 0.1 of OD_600nm_ with the indicated medium. After overnight incubation, OD_600nm_ values of the cultures were measured and plotted. When indicated, complete human serum (HS) was added to the culturing medium to 5% (v/v). The dashed line in A & B indicates 0.1 of OD_600nm_. For each tested condition, at least three biologically independent replicates were obtained and plotted with individual dots. For A & B, statistical analyses were performed with one-way ANOVA followed by Dunnett’s multiple comparison tests; For C & D, statistical analyses comparing between the growth with and without the supplement of human serum (i.e. between readings of green and orange of the same strain) were performed with the Mann-Whitney test with the *P*-values indicated in black above each box; statistical analyses comparing between different strains grown under the same condition were performed with the Kruskal-Wallis tests followed by Dunn’s multiple comparison tests against the WT values, *P*-values were indicated with the corresponding color of the culturing conditions inside the box. Nonsignificant *P*-value (β0.05) was indicated with “ns”; *, 0.01 < *P*-value < 0.05; **, 0.001 < *P*-value < 0.01; ***, 0.0001 < P-value < 0.001; ****, *P*-value < 0.0001.

**Fig 3:**
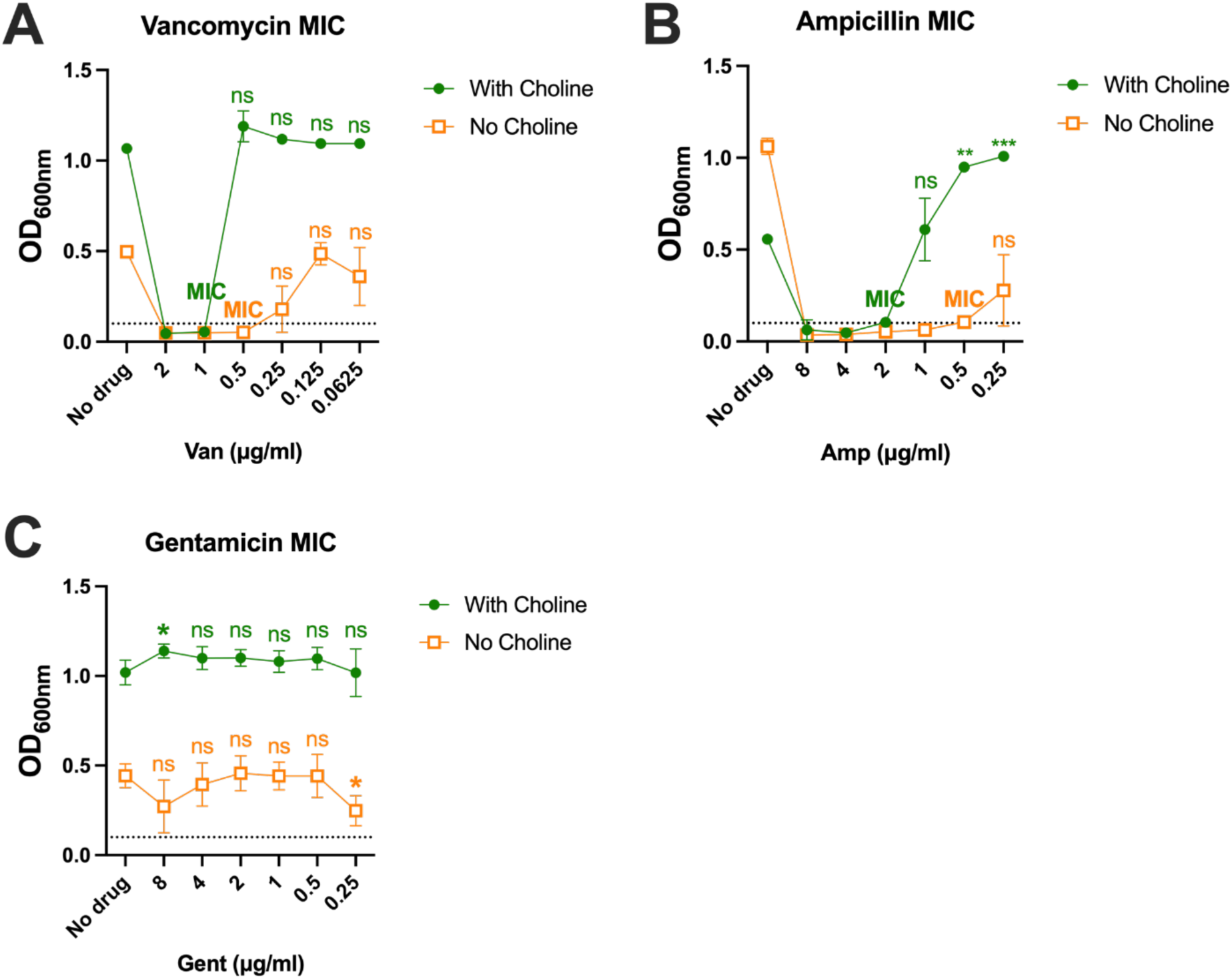
SM43 has increased susceptibility to vancomycin and ampicillin, but not gentamicin, when grown without choline. Stationary phase OD_600nm_ values of SM43 wildtype grown in either choline-supplemented chemically defined medium (CDM) (With Choline, green) or plain CDM (No Choline, orange) with the addition of vancomycin (A), ampicillin (B), or gentamicin (C) at the indicated concentrations. The dashed line indicates 0.1 of OD_600nm_, at which value the cultures were set with at the beginning of the incubation. For each tested condition, at least three biologically independent replicates were obtained. The minimum inhibition concentration (MIC) at each tested condition was determined by the lowest drug concentration at which a statistically significant growth inhibition was seen with no obvious bacterial growth. Statistical analyses were performed with one-way ANOVA followed by Dunnett’s multiple comparison tests controlled with the no drug values at the same tested condition. Nonsignificant *P*-values (β0.05) were indicated with “ns”; *, 0.01 < *P*-value < 0.05; **, 0.001 < *P*-value < 0.01; ***, 0.0001 < *P*-value < 0.001.

### Detection of Type IV LTA intermediates in SM43 provides experimental evidence about biosynthetic processes

In accordance with the observation that SM43 can grow without choline supplementation, a choline uptake-deficient mutant of SM43 was successfully generated. Specifically, the first gene of the operon encoding the choline uptake system, *licABC* (gene locus identifiers FD735_RS04490-04500), was replaced with *ermB* in SM43, producing the *ΔlicA* strain. Whole genome sequencing of the *ΔlicA* strain confirmed the deletion of *licA* and revealed additional mutations relative to the SM43 WT parent, including single nucleotide polymorphisms (SNPs) that resulted into ClpX^R80S^, PstB^L186fs^, and RplM^Y76C^ and in gene encoding phosphomevalonate kinase (FD735_RS08105, Ser303Cys) and ISL3 family transposase (FD735_RS08415, Leu194Ile) (Table S2).

Lipidomic analyses of *ΔlicA* by NPLC-ESI/MS detected several unexpected Type IV LTA biosynthetic intermediates (Fig. 1B-D) that are also observed in SM43 WT strains grown without choline, which serve as phenotypic controls for *ΔlicA*. Specifically, the accumulation of a single pseudopentasaccharide or TA repeating unit linked to MHDAG ([M-H]^-^ at *m/z* 1520.8; Fig. 1B&C, Fig S2A) and to undecaprenyl pyrophosphate (C_55_-PP) (Fig S2B) were detected. These intermediates, not surprisingly, lack P-Cho residues, and their accumulation suggests that P-Cho decorations are important for the polymerization of TA repeating units.

In addition, D-alanine (Ala) modifications were detected for both the C_55_-PP-linked pseudopentasaccharide ([M-2H]^2-^ at *m/z* 880.4 of Fig. 1D bottom panel) and MHDAG-linked pseudopentasaccharide ([M-H]^-^ at *m/z* 1591.8 of Fig. 1B), which does not support the model proposed by Fischer and colleagues (38–40) in which D-Ala is first attached to C_55_-P to form D-Ala-P-C_55_ which is then transported across the membrane to serve as the donor for D-Ala modification. In particular, the detection of D-Ala modification of C_55_-PP-linked pseudopentasaccharide refutes the proposed involvement of D-Ala-P-C_55_, a hypothesized lipid molecule that has not been detected by our highly sensitive lipidomic analysis. These experimental observations thus suggest the current model of Type IV LTA biosynthesis, which is largely based on bioinformatic analysis, needs to be revised and experimentally verified.

### Genes of the cpoA locus are responsible for synthesizing LTA glycolipid anchors

Mitis group streptococcal glycosyltransferases responsible for glycolipid biosynthesis were previously identified, with the function of *cpoA* verified (13, 30). Specifically, FD735_RS04120 (ortholog of *S. pneumoniae cpoA* (41)) is predicted to produce DHDAG, and FD735_RS04125 to produce MHDAG (Fig 4A). For convenience, FDR735_RS04125 is named in this study as *cpoC* (the gene name *cpoB* is already used for a cell division coordinator in *E. coli* (42)).

**Fig 4:**
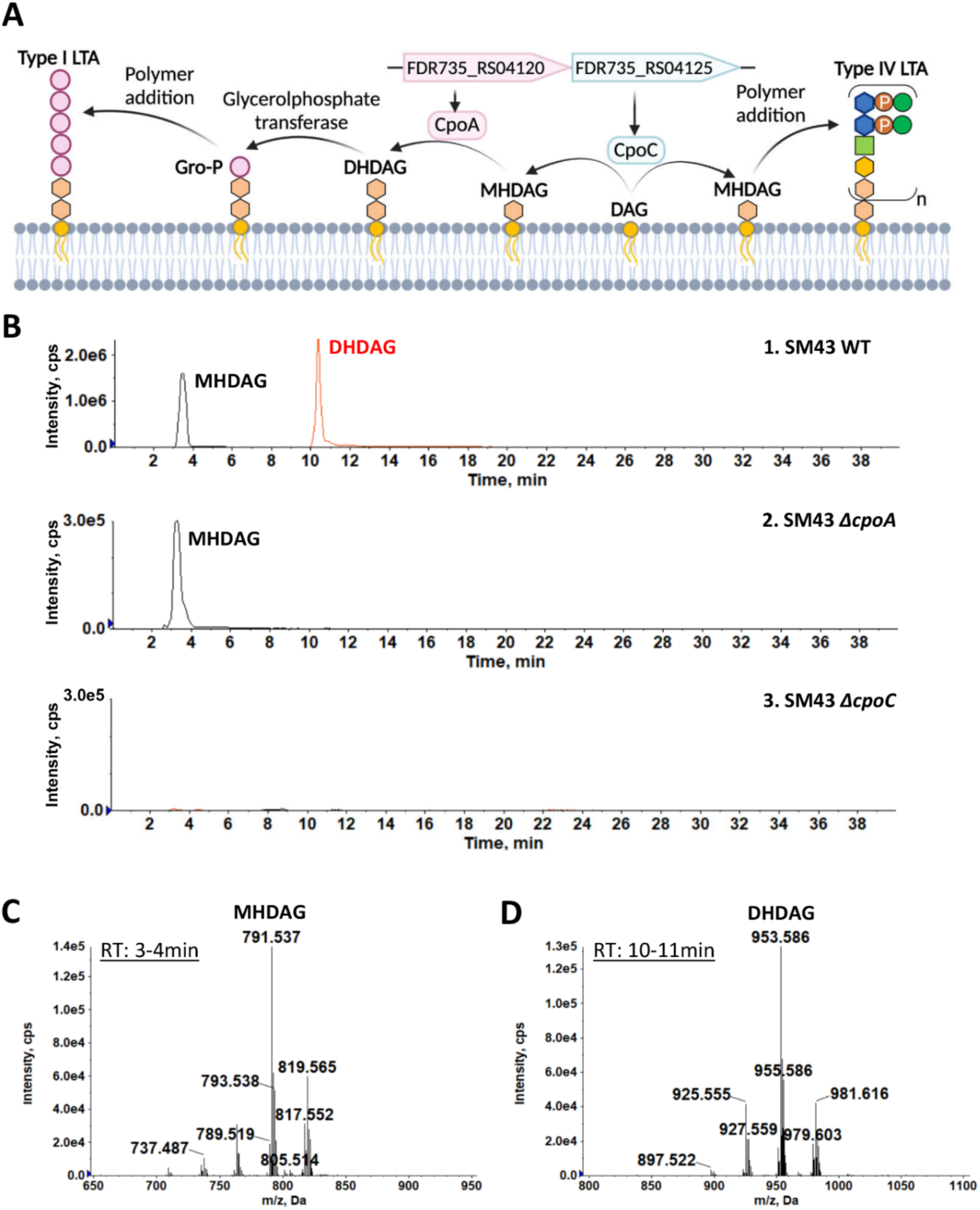
Glycosyltransferases CpoA and CpoC are responsible for *S. mitis* glycolipid biosynthesis. **A.** Diagram showing the gene locus of *cpoAC* in SM43 and predicted biosynthetic processes of glycolipid anchors and *S. mitis* LTAs. Chemical groups were presented with the same color coding as Fig 2A. **B-D.** Mass spectra of the [M+Cl]^-^ ions of monohexosyl-diacylglycerol (MHDAG) (retention time 3-4 minutes, most abundant *m/z* 791.537 for MHDAG (16:0/18:1) shown as C) and dihexosyl-diacylglycerol (DHDAG) (retention time 10-11 minutes, most abundant *m/z* 953.586 for DHDAG (16:0/18:1) shown as D) detected in SM43 WT (B1), *ΔcpoA* (B2), and *ΔcpoC* (B3)grown in TH broth. At least two biologically independent replicates were obtained for each tested strain at the indicated condition.

To verify the biosynthetic functions of *cpoA* and *cpoC*, individual *ermB* allelic replacement mutants were obtained. To mitigate polar effects, in each mutant, the coding region of either *cpoA* or *cpoC* was replaced by *ermB* coding region in the same direction. Additionally, genomic mutations in *ΔcpoA* and *ΔcpoC* strains were confirmed with whole genome sequencing (Table S2). It is worth noting that these strains each have different mutations in *glpQ*, a gene that encodes a glycerophosphodiester phosphodiesterase, which potentially can hydrolyze phosphatidylglycerol and produce glycerophosphate (43, 44). That the *ΔcpoA* and *ΔcpoC* strains have different, independently arising mutations (conferring Phe162Cys and Gly507Cys substitutions, respectively), suggests that mutation of this gene is associated with tolerating loss of cellular glycolipids. Additionally, SNPs in genes associated with peptidoglycan hydrolysis and cell replication were uniquely seen in the *ΔcpoC* mutant.

Lipidomic analyses were performed to verify the biosynthetic functions of *cpoA* and *cpoC*. As expected, DHDAG was not detected in the total lipids extracted from *ΔcpoA* (Fig 4B & Table 1) and neither DHDAG nor MHDAG was observed in *ΔcpoC* (Fig 4B). Additionally, GroP-linked glycolipids were missing in both *ΔcpoA* and *ΔcpoC* and no Type IV LTA intermediates were observed in both mutants either (Table 1), suggesting that DHDAG and MHDAG each respectively serve as the primary glycolipid anchor for GroP-DHDAG and Type IV LTA; and that DHDAG is also involved in the biosynthesis of Type IV LTA intermediates. Interestingly, an elevation of free fatty acids and production of another phosphatidic acid-derived glycolipid, phosphatidyl-*N*-acetylhexosamine (PAHN), is also observed in the *ΔcpoA* mutant (Fig S1). Increased production of PAHN was previously observed in *S. mitis* under stressful conditions, such as at the late stationary phase and in *S. mitis ΔcdsA*, a mutant that is deficient in producing phosphatidylglycerol, cardiolipin, and GroP-linked glycolipid, and is highly resistant to daptomycin (30, 45). The physiological functions and biosynthesis of PAHN remain unclear.

**Table 1:** Glycolipids and phospholipids detected in the wild-type and mutant strains of SM43.^a^.

| Genome type | PA | PG | CL | MHDAG | DHDAG | (Gro-P)-<br>DHDAG | LTA-IV<sup>b</sup> |
| --- | --- | --- | --- | --- | --- | --- | --- |
| WT | + | + | + | + | + | + | + |
| *ΔlicA* | + | + | + | + | + | + | + |
| *ΔcpoA* | + | + | + | + | - | - | -<sup>c</sup> |
| *ΔcpoC* | + | + | + | - | - | - | -<sup>c</sup> |

<sup>a</sup>Abbreviations and notations: +, detected; -, undetected; PA, phosphatidic acid; PG, phosphatidylglycerol; CL, cardiolipin; MHDAG, monohexosyl-diacylglycerol; DHDAG, dihexosyl-diacylglycerol; (Gro-P)-DHDAG, phosphoglycerol-DHDAG.
<sup>b</sup>Refers to Type IV LTA intermediates.
<sup>c</sup>No detection of Type IV LTA intermediates in samples grown with TH broth.

### Glycolipids are required for optimum growth of SM43

All of the reported mutants (i.e. *ΔlicA*, *ΔcpoA*, *ΔcpoC*) show significant growth deficiency compared to the WT strain (Fig 2B) in both laboratory undefined (Todd Hewitt broth) and chemically defined medium (CDM) (Fig 2C), except in CDM without choline (Fig 2D), under which condition no significant growth difference was observed between WT and *ΔlicA*. The results demonstrate that DHDAG and MHDAG are required for optimum growth. Interestingly, when grown with TH broth, *ΔcpoC* showed no significant growth difference as compared to the *ΔlicA* strain (Fig 2B) with both strains demonstrating significant growth deficiency compared to *ΔcpoA*, highlighting the roles of glycolipids and properly anchored LTA in the growth of SM43.

To evaluate the effects of glycolipid species in bacterial growth under host-associated conditions, such as inside the gingival pockets and bloodstream, human serum was added to the culture media. Similar to what has been previously observed (30, 46), supplementation of human serum significantly promoted the growth of the SM43 WT strain irrespective of the presence or absence of choline supplementation in the culture media (Fig 2C & D). Similar growth-promoting effects were also observed in glycolipid-deficient strains when choline supplementation was present (Fig 2C), but not when choline was absent (Fig 2D). Additionally, such growth promotion effects were not seen in *ΔlicA* (Fig 2C) no matter whether choline was added or not. Results indicate the crucial (but non-essential) role of choline in the optimum growth of SM43 and suggest that the presence of 5% serum cannot fully compensate for the lack of supplemental choline.

### Glycolipid-deficient SM43 is attenuated in a murine sepsis model

While there are well-established models for *S. pneumoniae* (47–49), *in vivo* bacteremia models of *S. oralis* or *S. mitis* infections remain scarce (50). Therefore, we first established and characterized a murine sepsis model for SM43 to evaluate its virulence. In this case, we are comparing the virulence between the SM43 WT and mutant strains to establish the contributions of glycolipids and the SM43 cell surface to virulence and host response.

To establish a sepsis model, an infectious dose-response curve was generated in C57BL/6J male and female mice, with tail-vein injections of mid-exponential phase SM43 (Fig. 6A). Doses of 2.5 x 10^9^ CFU/mouse and 1.0 x 10^9^ CFU/mouse were found to be 100% lethal within 24 hours post-infection, while doses of 2.5 x 10^8^ CFU/mouse and 1.0 x 10^8^ CFU/mouse resulted in 100% survival over 3 days post-infection (Fig 6A). Following a stepwise pattern, partial lethality (60%) was observed with a dose of 5 x 10^8^ CFU/mouse (Fig 6A). The lowest fully lethal dose, 1.0 x 10^9^ CFU/mouse, was then selected for further virulence analysis. Using this model, the virulence of SM43 WT was contrasted with the Δ*cpoA* and Δ*cpoC* mutants. The Δ*cpoC* mutant exhibited a marked reduction in virulence, with lethality dropping from 100% in SM43 WT to 40% in the mutant (Fig 6B).

### Functions of glycolipids in protecting against cell wall-targeting antibiotics

Previous studies identified decreased susceptibility to β-lactams in pneumococcal *cpoA* mutants (34). As indicated by Table 2, the SM43 mutants generated in this study have increased susceptibility toward all tested antibiotics. These results align with the observation that the absence of choline in the culture medium leads to increased susceptibility to cell surface-targeting antibiotics, such as ampicillin and vancomycin, but also suggest further functions of the glycolipids in modulating membrane permeability, as the susceptibility towards the ribosome-targeting antibiotic gentamicin also increased. Visualization of SM43 WT, *ΔcpoA*, and *ΔcpoC* cells via sectional transmission electron microscopy (TEM) revealed increases in the thickness of the cell surface structure in both mutants (Fig 5), but no obvious differences in cell shape or arrangement.

**Table 2:** Antibiotic minimum inhibitory concentrations (MIC, median and range in μg/ml, n ≥ 3) of *Streptococcus* sp. SM43 wildtype (WT) and mutants measured with E-test strips.

| Antibiotic | WT | *ΔcpoA* | *ΔcpoC* | *ΔlicA* |
| --- | --- | --- | --- | --- |
| Daptomycin | 0.38 (0.380 – 0.75) | 0.047 (0.047 – 0.064) | < 0.016 | 0.125 (0.125 – 0.19) |
|  | **Fold decrease from WT:** | **8.09** | **>23.75** | **3.04** |
| Gentamicin | 4.0 (4.0 – 6.0) | 3.0 (3.0 – 4.0) | 0.016 (0.016 – 0.016) | 1.0 (1.0 – 1.0) |
|  | **Fold decrease from WT:** | **1.33** | **250.00** | **4.00** |
| Vancomycin | 0.5 (0.5 – 0.75) | 0.19 (0.05 – 0.5) | 0.19 (0.19 – 0.38) | 0.19 (0.125 – 0.25) |
|  | **Fold decrease from WT:** | **2.63** | **2.63** | **2.63** |
| Fosfomycin | 24 (24 – 32) | 12 (8 – 16) | 2 (1.5 – 4) | 0.250 (0.19 - 0.25) |
|  | **Fold decrease from WT:** | **2.00** | **12.00** | **96.00** |
| Ampicillin | 0.25 (0.25 – 0.38) | 0.25 (0.19 – 0.25) | 0.094 (0.084 – 0.125) | < 0.016 |
|  | **Fold decrease from WT:** | **1.00** | **2.66** | **15.63** |

**Fig 5:**
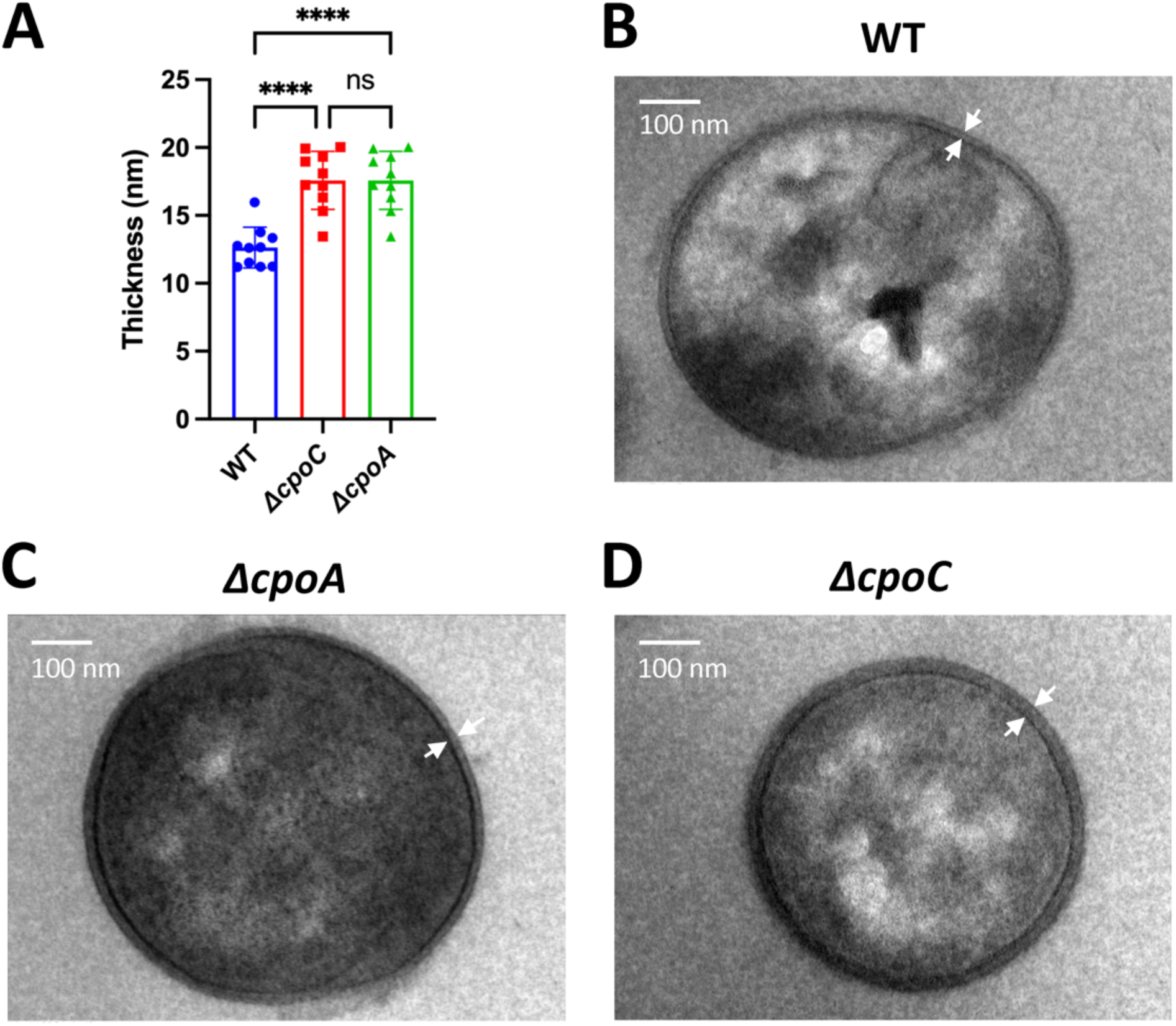
Mutation of glycosyltransferases results in thickened cell surface structure in SM43. **A**. Plot of the thickness of the cell surface structure (indicated by the arrows in panels B-D) of SM43 WT and mutants. **B-D**. Transmission electron microscope image of SM43 WT and mutant cells. Images were taken at 25,000ξ total magnification. Scale bars indicate 100nm. Statistical analyses were performed with one-way ANOVA followed by Dunnett’s T3 multiple comparisons test. A nonsignificant *P*-value (β0.05) is indicated with “ns”; **** stands for P-value < 0.0001. For panel B, statistical analyses were performed with one-way ANOVA followed by Tukey’s multiple comparisons test with *P*-values listed in Table S3.

**Fig 6.**
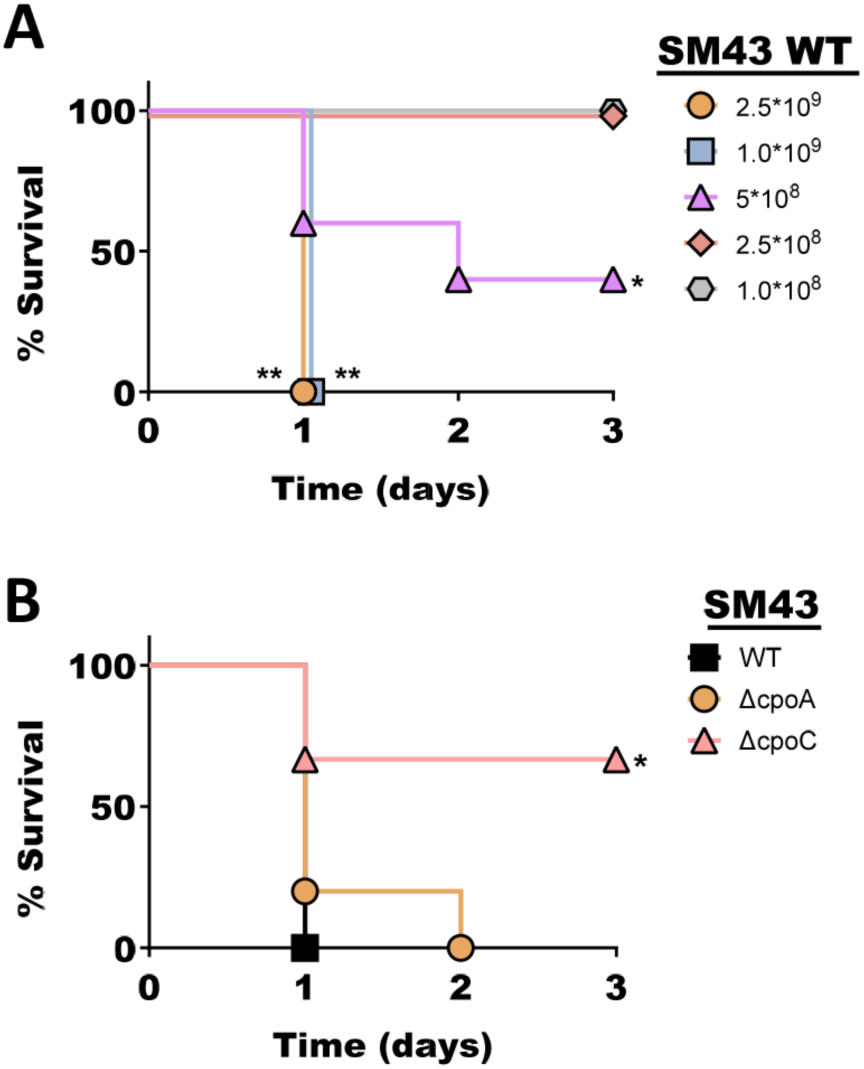
Murine sepsis model for SM43 demonstrates virulence defect in *ΔcpoC*. **A** murine sepsis model was established using SM43, with C57BL/6J mice (n=5 per groups) infected via the tail vein injections. A. Mice were injected with wildtype (WT) SM43 at the indicated doses (CFU/mouse) to assess dose-dependent lethality. B. Mice were injected with 1×10^9^ CFU/mouse of either SM43 WT, Δ*cpoA,* or Δ*cpoC* strain to evaluate virulence defects in vivo. Statistical significance was determined using Kaplan-Meier survival analysis, comparing (A) survival rates at 1×10^8^ CFU/mouse, or (B) SM43 WT versus mutant strains, with significance denoted as “*” for 0.01 < *P*-value < 0.05, “**” for 0.001 < *P*-vlaue < 0.01.

## Discussion

In this work, we verified that SM43 glycosyltransferases CpoA and CpoC are each responsible for producing DHDAG and MHDAG, which correspondingly serve as the lipid anchor of GroP-DHDAG and Type IV LTA. Additionally, we detected Type IV LTA intermediates that contradict the predicted biosynthetic process of cytoplasmic polymerization of TA units, but instead suggest the cross-membrane translocation before full assembly of the TA polymers (Fig 3). Interestingly, the *S. pneumoniae* TA polymerases (TarQP) are predicted to be extracellular enzymes, which also suggests that the polymerization of TA units happens on the outer leaflet of the membrane (51, 52). However, SM43 has no homolog to either *tarQ* or *tarP* (Table S1). New extracellular genes involved in the Type IV LTA biosynthetic process still await identification.

It is worth noting that though SM43 does not require choline for survival, the presence of a sufficient amount of choline is necessary for optimum growth. When the cells are deficient in choline uptake, such as the *ΔlicA* strain, the growth-promoting effect of human serum is abolished. Deficiency in producing either CpoA or CpoC also renders the cells nonresponsive to the growth-promoting effects of human serum when choline is not supplemented in the medium. According to the Human Metabolome Database (53), human blood contains 6.0-27.5 μM choline at normal conditions. We added human serum to 5% in the culturing media, which resulted in a final choline concentration of no more than 1.4 μM, which is less than 0.3% of the choline (final concentration at 500 μM) that we supplemented in the chemically defined medium; and thus, cannot adequately compensate for the lack of choline in the medium. At certain conditions, the human blood choline concentration will increase up to > 140 μM (53) (such as in newborns and pregnant women), which would support the optimum growth of MGS and potentially make people with this condition more susceptible to MGS infections. This hypothesis can be verified with future epidemiological data.

To further evaluate the roles of *S. mitis* glycolipids and associated cell surface components in pathogenesis, a murine sepsis model was developed for virulence evaluation. Interestingly, even though no significant growth difference is observed between the WT and *ΔcpoC* grown in choline-supplemented CDM with 5% human serum, *ΔcpoC* is significantly attenuated in lethality, making *cpoC* a potential target for therapeutic design of treatment.

Aside from supporting optimum growth and complete virulence inside of the host, Type IV LTA and glycolipids also seem important for maintaining membrane stability and selective permeability, as the mutants have increased susceptibilities towards antimicrobials targeting cell surface structures and antimicrobials targeting cytoplasmic biological processes. Additionally, the fact that SM61, which also produces Type I LTA, cannot grow without choline, suggests that Type I LTA cannot compensate for the cellular functions of Type IV LTA, leaving the functions of Type I LTA in MGS unknown. It was previously observed that the absence of choline, which hinders Type IV LTA production, leads to abnormal cell shapes, chain formation, and smaller colony size in *S. pneumoniae* (25, 37). Our TEM images suggest less dramatic effects of missing glycolipids in SM43, which leads to the question of whether the transfer of the TA polymer to its corresponding glycolipid anchor is necessary for the performance of its physiological functions, at least in SM43. Moreover, elongated chains were seen for some mutant strains generated in this study, but the phenotype was inconsistent, with varied cellular morphologies among different cultivation times and media (data not shown). It is possible that the expression of LTA is a dynamic process that changes with the growth phase and is modulated by environmental conditions, and that the functions of LTA also vary accordingly. One example supporting this idea is the transfer of TA polymers from lipid anchors to peptidoglycan in *S. pneumoniae* grown to the stationary phase, which leads to autolysis (54).

One major downside of this study is that genomic complementation of the deleted genes could not be constructed. Complementary plasmids carrying the deleted genes (*licABC*, *cpoA*, and *cpoC*) were generated but failed in transformation, using either the natural competence-induced transformation or electroporation methods described previously (30, 55). It has been reported that cell surface component biosynthetic genes are associated with competence and autolysis (56), which could be the cause of transformation failures. In all reported deletion mutants, mutation of other genes was observed. Interestingly, SNPs in *clpX* are seen in *ΔlicA* mutant. In *S. aureus*, mutation of *clpX* permitted survival without Type I LTA, results suggesting an epistatic interaction between LTA and ClpX (57). Similar epistatis between ClpX and Type IV LTA is potential in MGS. ClpX is the ATPase and substrate recognition component of the protease complex ClpXP, which is essential for growth and highly conserved in bacteria, mitochondria, and chloroplasts and involved in various cellular processes, including DNA damage repair and bacterial virulence (58–60). In *S. pneumoniae*, ClpX also regulates the development of competence (51), affecting the successful rate of transformation.

Despite the long history of Type IV LTA identification (61), there are still many unanswered questions regarding its detailed biosynthetic processes and its functions. For example, except *tacF*, no other gene with putative flippase activity was identified in MGS; yet, based on the predicted enzyme locations, glycolipid biosynthesis happens at the cytoplasmic side, indicating the existence of an uncharacterized flippase. Additionally, despite the severe growth deficiency, the *ΔcpoC* mutant was successfully generated and the mutant devoid of major glycolipids has a normal cell shape, leading to interesting questions on the minimal lipid requirement of Gram-positive bacterial membranes. Answers to these questions will help us obtain a fundamental understanding of bacterial physiology.

## Materials and methods

### Bacterial strains and culturing conditions

*S. mitis* NCTC12261^T^ and the infective endocarditis isolate *Streptococcus* sp. 1643 (29) and its mutants were all grown at 37°C with 5% CO_2_ with either Todd-Hewitt medium (TH medium; BD Biosciences), chemically defined media (CDM), or Mueller-Hinton medium (MH medium, BD Biosciences). CDM is prepared as described before (62). When noted, the final concentrations of the added compounds were as follows: 0.5 mM choline; 5% (v/v) of complete human serum (Sigma-Aldrich; H6914); erythromycin, 20 μg/ml in streptococci and 50 μg/ml in *E. coli*.

### Homolog identification

Gene orthologs were identified by using the BLASTp function against the nonredundant protein database of either *Streptococcus sp*. 1643 (SM43, taxid: 2576376) or *S. mitis* NCTC 12261^T^ (SM61, taxid: 246201) with a query coverage ≥ 93% and E-value ≤ 10^-35^ (63). Previously predicted lists of Type IV LTA biosynthetic genes in *S. pneumoniae* R6 and *S. oralis* Uo5 were used as references (13, 18). Orthologs of Type IV LTA biosynthetic genes in SM43 and SM61 were listed in Table S1.

### Deletion mutant generation

Deletion of *licA* (FD735_RS04490), *cpoA* (FD735_RS04120), and *cpoC* (FD735_RS04125) in SM43 was conducted as described before (30). Specifically, a 5kb DNA fragment that sequentially contains a 2kb fragment upstream of the target gene, a 1kb fragment containing *ermB* in the needed direction, and a 2kb fragment downstream of the target gene was generated via overlapping PCR followed by cleaning with the PCR cleaning kit (ThermoFisher). Transformation of the 5kb amplicon into SM43 was performed as described previously (64), and successful transformants were selected with erythromycin. Mutant candidates were confirmed with Sanger sequencing of the mutated region performed by the Genome Center at The University of Texas at Dallas (Richardson, TX), and then sent for whole genome sequencing to identify the existence of any other mutations. Specifically, sequencing of the *ΔlicA* mutant was performed by the Genome Center at The University of Texas at Dallas (Richardson, TX, USA); whole genome sequencing of SM43 isogenic WT and mutants *ΔcpoA* and *ΔcpoC* were performed by the SeqCenter (Pittsburgh, PA, USA). Mapping of the sequencing reads and detection of the SNPs were performed with CLC Genomics Workbench (version 20; Qiagen). Whole genome reference of SM43 (NZ_CP040231; taxid: 2576376) was downloaded from NCBI database. SNPs shared between the isogenic WT and deletion mutants were excluded from analyses. All primers used in this study are listed in Table 3.

**Table 3:**
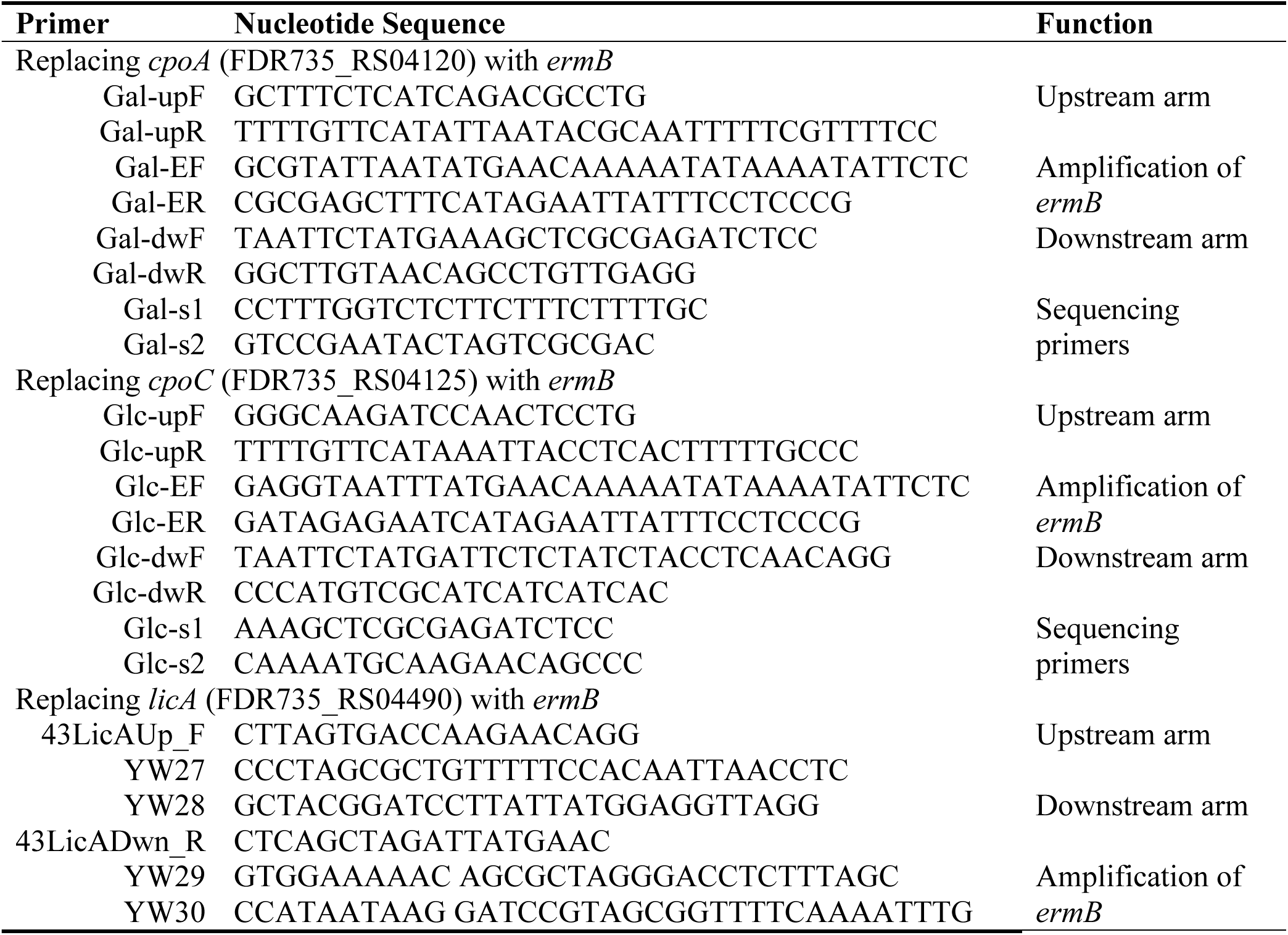
Primers used in this research.

### Measurement of bacterial antibiotic susceptibility

Antibiotic susceptibility of SM43 WT strain grown with and without the presence of choline was measured via a broth microdilution method modified from the previous description (65). Specifically, a two-fold serial dilution of the antibiotic was made with CDM either with or without choline in a 96-well microtiter plate starting from row C to H, leaving 100 μl of liquid in each well. 200 μl and 100 μl of the same medium in each column were added to wells in rows A and B correspondingly. Bacterial cultures at 0.2 of OD_600nm_ were added to all wells in row B-H in a 1:1 volume ratio. Prepared plates were incubated overnight before measurement of the OD_600nm_ readings. According to the definition of the minimum inhibitory concentration (MIC) provided by the Clinical and Laboratory Standards Institute (CLSI) guidelines (66), SM43 MICs were determined by the lowest antibiotic concentrations that resulted in < 0.1 OD_600nm_ value after subtracting the average value of blank (plain medium). To prepare bacterial cultures used for the plate, SM43 cells were pelleted (centrifuge at 15,000 g for 6 minutes) from single-colony overnight cultures grown in CDM with choline, followed by resuspension in CDM without choline into 0.1 of OD_600nm_. After overnight incubation of the resuspended cells, bacterial cultures were diluted into 0.2 of OD_600nm_ in CDM either with or without choline supplement, incubated for 1 hour, and then added into the plate as described above. The highest testing concentrations of the antibiotics were calculated as 4 times the previously reported MICs (29, 66, 67): 8 μg/ml ampicillin, 2 μg/ml vancomycin.

Antibiotic susceptibility on agar plates was measured with E-test strips per the manufacturer instructions. Specifically, E-test strips of daptomycin, gentamicin, vancomycin, and fosfomycin were purchased from BioMerieux; E-test strips for ampicillin were purchased from Liofilchem. Bacteria single colony was grown in either TH or MH broth overnight. If grown with TH broth, bacterial cells were pelleted and resuspended in MH broth to 0.1 of OD_600nm_ and incubated overnight. Bacteria grown in MH broth were spread on MH plates with sterile cotton-tipped applicators, followed by 15-20 min air-dried under the biosafety cabinet before application of the E-test strips by sterilized metal forceps. After 24 hours of incubation, the minimum inhibitory concentration (MIC) was determined by the intersection of the zone of inhibition with the E-test strip. For each tested condition, at least three biologically independent replicates were obtained.

### Lipidomic analyses

Lipidomic analyses were performed exactly as previously described (30, 68). Total lipid samples were extracted with the modified acidic Bligh-Dyer method from a minimum of 5ml bacterial cultures grown under the indicated conditions (30). For each tested condition, at least two biologically independent replicates were obtained for verification.

### Imaging of bacterial cells

TEM images were taken by the Imaging Core Facility at the Oklahoma Medical Research (Oklahoma City, OK). Bacterial cells were pelleted from 5ml overnight cultures grown in TH broth, washed with PBS, and treated with 2.5% glutaraldehyde prepared in PBS (pH 6.5-7.4) at room temperature for 1 hour before being sent for imaging. For each tested train, 10 individual cells were imaged. ImageJ was used to measure the thickness of the cell surface structure at 10 randomly selected locations for each imaged cell.

### Murine sepsis model

SM43 WT and its mutants, *ΔcpoA* and *ΔcpoC*, were grown overnight at 37°C in 5% CO2 in TH broth, followed by dilution into 0.1 of OD_600nm_ with fresh TH broth and resumed incubation. When the OD_600nm_ values reached 0.3 to 0.6, bacterial cells were harvested with centrifugation at 5,000 g for 10 minutes. Cell pellets were resuspended with sterile 1× PBS at a 1/10 volume of the culturing media, followed by centrifugation at 10,000 g for 5 min. Pelleted cells were resuspended to the desired concentration (CFU/ml) in 1× DPBS (Dulbecco’s) for injection.

In the sepsis model, 7-8 week-old C57BL/6J mice (Jackson Laboratories) were placed in a tail vein restrainer (Braintree Scientific). 100 µL of the prepared bacterial suspension was administered via the lateral tail vein using an insulin syringe (BD Medical). Equal numbers of male and female mice were included in the study, and their survival was monitored for 3 days post-infection. Post-infection/treatment, mice were monitored every 6 to 12 hours for any signs of severe lethargy or agitation, moribund appearance, or failure to right oneself after 5 seconds. Animals with these symptoms were humanely sacrificed by CO_2_ exposure followed by cervical dislocation and the time of death was recorded. All animal protocols were approved by the UTD IACUC.

### Data availability

Raw data of whole genome sequencing have been uploaded to the NCBI SRA database with ProjectID PRJNA1005251 and BioSample accession numbers SAMN36977669, 71, 72, &74.

## Acknowledgements

Fig 1A&E and Fig 4A are created with Biorender.com.

We thank the Genome Center at The University of Texas at Dallas and the Imaging Core Facility at the Oklahoma Medical Research for their services to support our research.

This work was supported by grants and R01AI148366 from the National Institutes of Health to K.L.P. and Z.G, the Cecil H. and Ida Green Chair in Systems Biology Science to K.P, and University of Texas at Dallas start-up funds to N.D.

---

**Fig S1:**
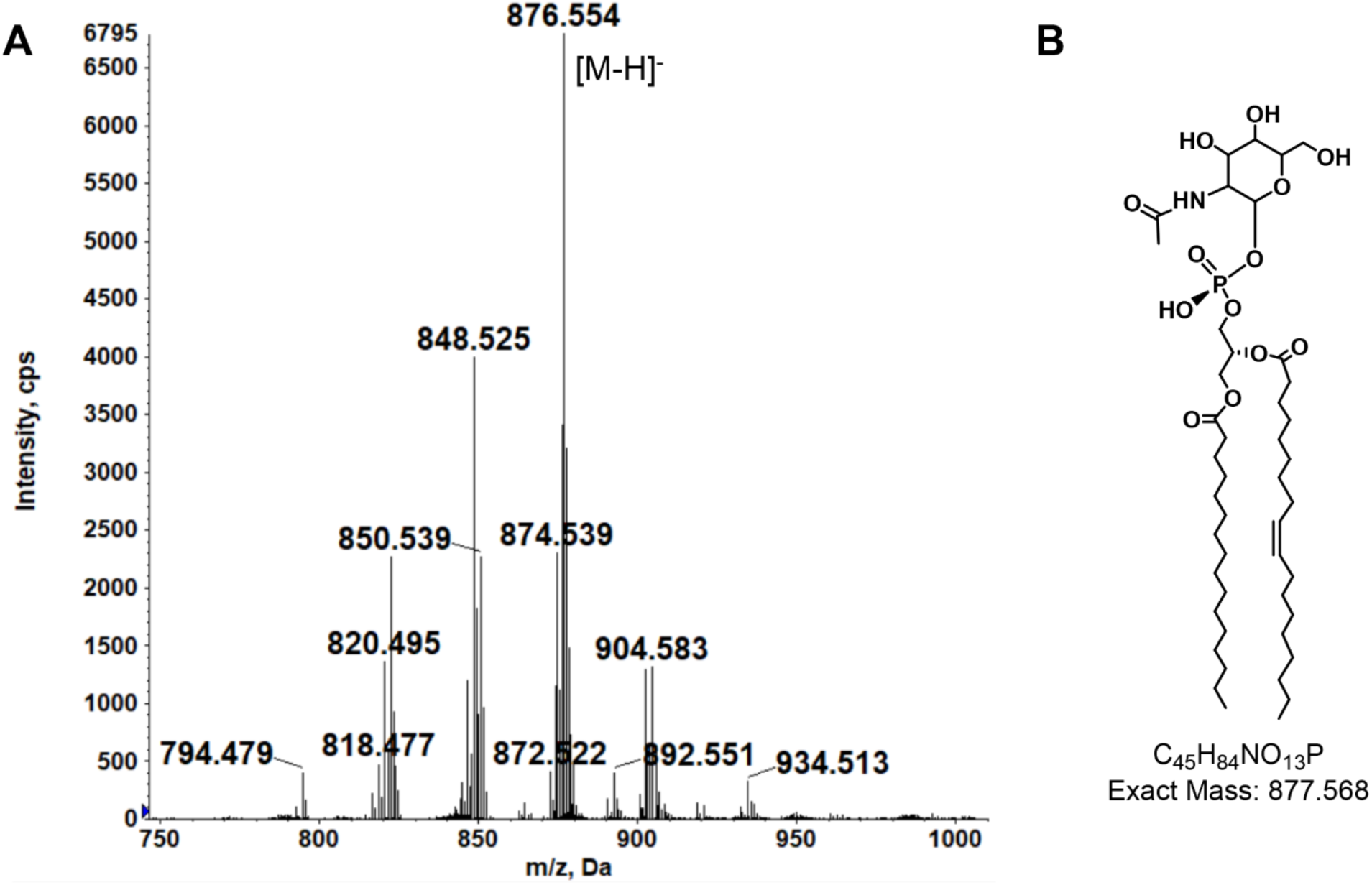
MS detection of phosphatidyl N-acetylhexosamine (PAHN) in the Δ*cpoA* mutant. A. Negative ion mass spectrum of [M-H]^-^ ion species of PAHN. B. PAHN (16:0/18:1) chemical structure.

**Fig S2:**
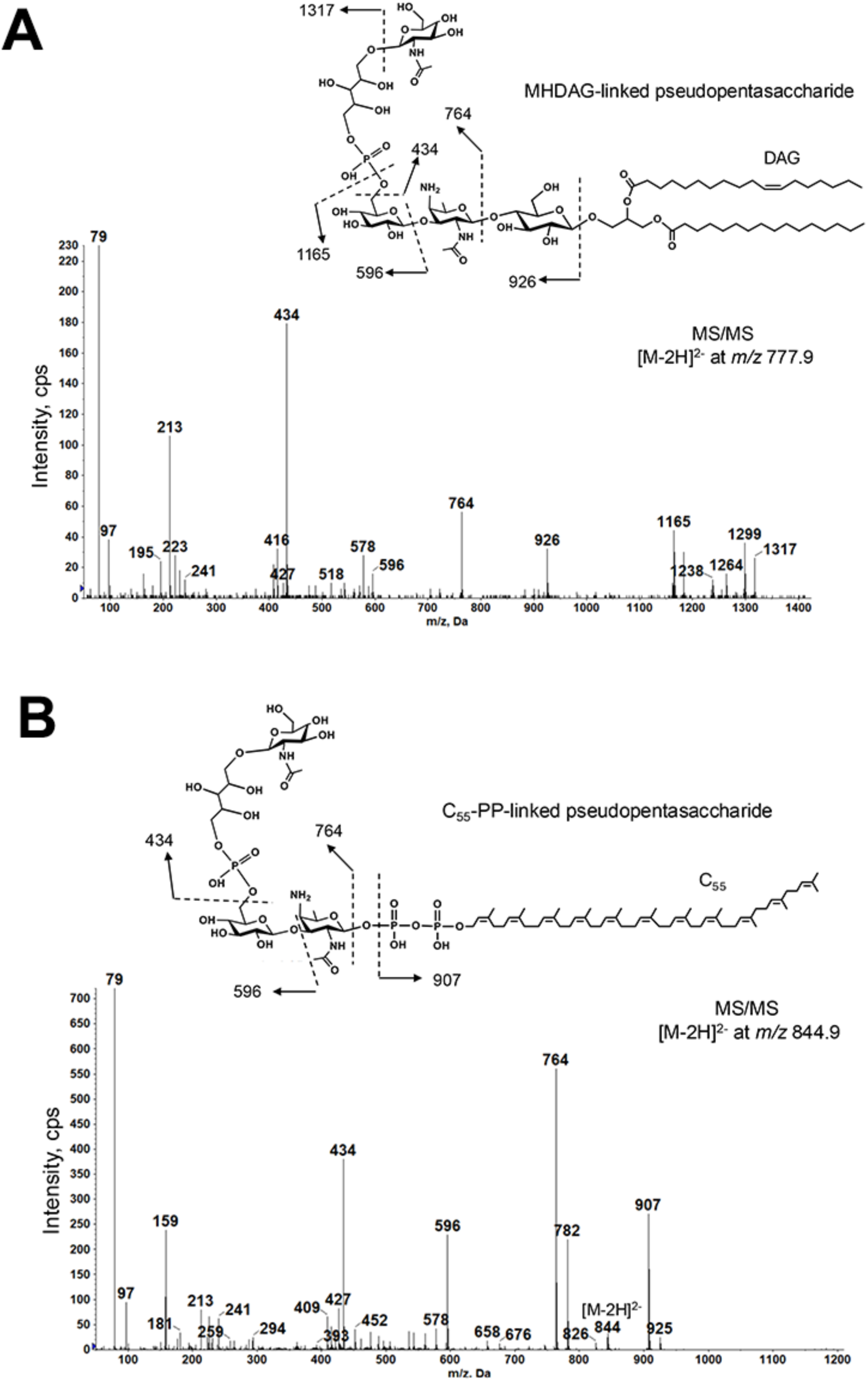
MS/MS structural confirmation of undecaprenyl pyrophosphate (C_55_-PP)-linked pseudopentasaccharide (A) and monohexosyl-diacylglycerol (MHDAG) -linked pseudopentasaccharide (B).

